# Comprehensive Cell Type Specific Transcriptomics of the Human Kidney

**DOI:** 10.1101/238063

**Authors:** V Sivakamasundari, Mohan Bolisetty, Santhosh Sivajothi, Shannon Bessonett, Diane Ruan, Paul Robson

**Author notes:** Equal contribution (V Sivakamasundari, Mohan Bolisetty, Santhosh Sivajothi). Corresponding author: Paul Robson, The Jackson Laboratory for Genomic Medicine, 10 Discovery Drive, Farmington, CT, USA 06032.

## Abstract

The human kidney is a complex organ composed of specialized cell types. To better define this cellular complexity, we profiled the individual transcriptomes of 22,469 normal human kidney cells, identifying 27 cell types. We describe three distinct endothelial cell populations, a novel subset of intercalated cells, interstitial macrophage and dendritic cells, and identify numerous novel cell-type-specific markers, many validated using imaging mass cytometry and immunohistochemistry. Receptor-ligand analysis revealed previously unknown intercalated-endothelial and intercalated-distal nephron interactions, suggesting a role in maintenance of vascular integrity and intercalated cell survival. Notably, kidney disease-associated genes were largely expressed in proximal tubules, podocytes, endothelial and myeloid cells, highlighting an underappreciated role for endothelial cells in kidney pathologies. Our analysis also provides a resource of cell type enriched markers, solute carriers, channels and lncRNAs. In summary, this cell-type-specific transcriptome resource provides the foundation for a comprehensive understanding of kidney function and dysfunction at single cell resolution.

---

The kidney is a complex organ composed of numerous cell types performing essential functions of filtering metabolic waste, balancing blood electrolytes and maintaining blood pressure. Kidney dysfunction has a significant impact on human health - chronic kidney disease (CKD) affects 10% of the worldwide population and 14% within the US resulting in 1.1 million deaths every year and costing nearly $100 billion in 2015^1,2^. Adult kidney is composed of hundreds of thousands of nephrons, which allow intricate exchange of materials between the blood and urine compartments. Decades of research using lineage-tracing in animal models and histology on human biopsies have extensively described the kidney architecture and functions^3^.

The nephron is composed of multiple distinct compartments: glomerulus, proximal tubule, distal tubule, connecting tubule and collecting duct. These segments perform different aspects of the filtration process, beginning with the glomerulus where primary urine is first formed. At least 20 distinct cell types have been defined in the literature with 4 distinct cell types in the glomerulus alone - endothelial cells of the glomerular tuft, podocytes, mesangial cells and parietal epithelial cells forming the Bowman’s capsule. Primary urine within the Bowman’s space then flows into the proximal tubules which are lined with epithelial cells that alter the repertoire of proteins expressed as distance from the glomeruli increases. Epithelial cells of the distal tubule segments are well-characterized by molecules marking distinct positions along its length. However, once reaching the connecting tubules and collecting duct there is distinct heterogeneity within the epithelium with principal cells interspersed with acid-base regulating intercalated cells. Other cell types that form the kidney interstitium are fenestrated and non-fenestrated endothelial cells of the peritubular capillaries, vascular smooth muscle cells, pericytes, fibroblasts and resident immune cells^4–8^.

A comprehensive molecular and spatial knowledge of the human kidney – a cell atlas - is currently lacking. Single-cell technologies and multi-parameter spatial imaging techniques now enable the construction of such an atlas^9^. Kidney pathology is not restricted to a single cell type or renal structure - glomerular and tubular pathophysiology can be mediated through epithelial, endothelial, mesenchymal, and/or immune cell dysfunction^10^. Hence, a complete transcriptome of each cell type within the kidney will not only inform on their function within the normal kidney but aid in understanding the origin of various kidney pathologies^10,11^.

In this study, we profiled 22,469 cells from the normal human kidney by droplet-based single cell transcriptomics, identifying 27 cell types. We describe new markers for known cell types and define new subsets within the intercalated and endothelial cell populations. We also describe interactions between different cell types in the kidney revealing a potential role for endothelial cells in kidney pathologies. In addition, we utilize imaging mass cytometry (IMC) and traditional immunohistochemistry (IHC) to provide spatial context to cell types and associated markers. We also resolve the expression of genes associated with kidney pathologies which point to the primary cell type(s) of dysfunction in these diseases.

## Results

To investigate the cellular composition in normal kidney, we adopted a high throughput single-cell transcriptomic approach. Normal kidney resection samples were enzymatically dissociated into a single cell suspension, then sorted for viable cells, before profiling using the Chromium™ system (**Supplementary Fig. 1**). We validated the cell type specific expression and spatial distribution of markers using IMC and IHC on formalin-fixed paraffin-embedded (FFPE) sections of normal kidney tissues (**Supplementary Fig. 7**).

We initially optimized tissue dissociation and cell profiling utilizing the first generation of Chromium™ chemistry, generating 3,128 single cell transcriptomes (**Supplementary Fig. 8**). We subsequently profiled two additional kidney resections, one male and one female, with version 2 Chromium™ chemistry and used this later data for comprehensive cell type identification. This represented a data set of 19,341 cells with an average of 1,018 genes and 2,462 median UMIs (Unique Molecular Identifiers) detected per cell (**Supplementary Fig. 2**). We employed unbiased clustering and dimensionality reduction methods to identify cell types. For cell typing, markers for each cluster were determined using Area Under Receiver Operating Characteristic (AUROC) analysis with genes greater than 2-fold difference in a cluster relative to all cells (Fig. 1, **Supplementary Fig. 4 and Supplementary Table 1**). We identified 8 distinct clusters from these 19,341 cells, with proximal tubule (PT) epithelial cells as the largest cluster (n=16,348; 84.5%) the predominant cell type in the kidney cortex and medulla (Fig. 1a).

**Figure 1:**
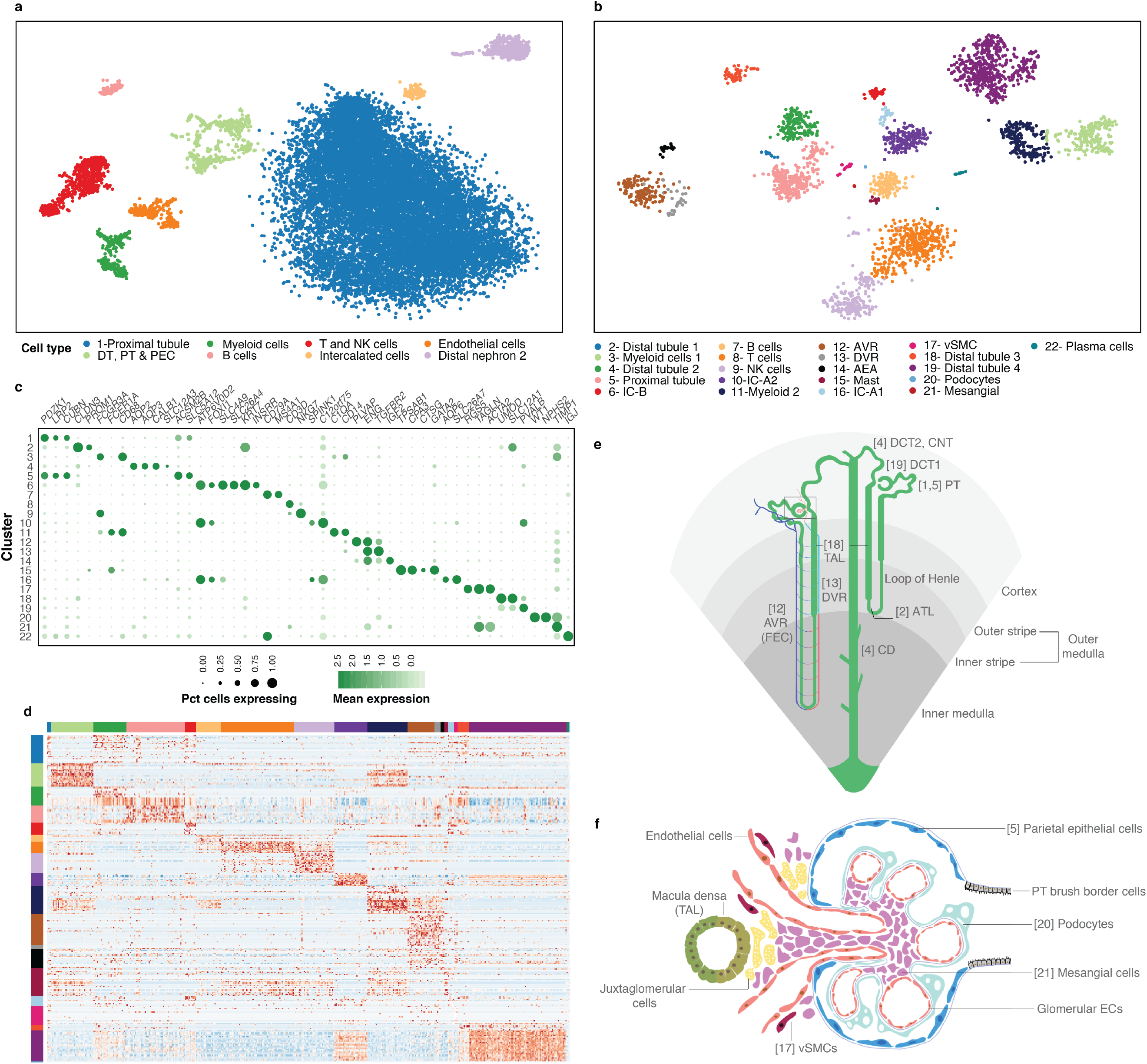
Human kidney cell types delineated by single cell transcriptome. (a) Unsupervised clustering of combined non-immune and immune cell types of human kidney represented as a tSNE plot. Different cell type clusters are color coded. (b) Cells were re-clustered (unsupervised) after removal of proximal tubule cell cluster shown in (a), revealing finer subsets of various non-immune and immune cell types, totaling 22 clusters. Different cell type clusters are color coded. (c) Bubble plot showing selected cell type specific markers across all 22 clusters. Size of dots represents percentage of cells expressing a particular maker and intensity of color indicates level of mean expression. Legends are shown below. (d) Heatmap showing scaled expression (log CPM) of discriminative marker gene sets across all 22 clusters, with cells as columns and rows as genes. Color scheme represents Z-score distribution with −3 (blue) to 3 (dark orange). (e) Illustration of cortical and juxtaglomerular nephrons and the various tubular segments of the kidney nephron. Spatial localization of nephrons across cortical, inner and outer medullary regions is demarcated. (f) Illustration of the glomerular region boxed in (e), showing various cell types present within the glomerulus and the adjacent macula densa segment of the thick ascending limb. Illustrations are not drawn to scale and corresponding cluster numbers are shown in [ ]. Abbreviations: NK – natural killer; IC-A – intercalated cell type A; IC-B – intercalated cell type B; EC – endothelial cell; Pct – percentage; PT – proximal tubule; DT – distal tubule; ATL – ascending thin limb; TAL – thick ascending limb; CD – collecting duct; AVR – ascending vasa recta; DVR – descending vasa recta; FEC – fenestrated endothelial cells; vSMCs – vascular smooth muscle cells; AEA – Afferent and efferent arterioles; PEC – Parietal epithelial cells; DCT – Distal convoluted tubule.

We further resolved the smaller subsets and rarer cell types by re-clustering the remaining 2,993 non-PT cells (Fig. 1b). This identified 21 clusters, including a cluster (cluster 5) of 311 PT-like cells that lay outside the original PT cluster 1 (Fig. 1b). We then integrated the original PT cluster with these 21 clusters to derive 22 clusters for downstream analyses (Fig. 1a-d). Subsequent cell type identification of each cluster was informed by markers from published studies and assisted by an analysis of the entire repertoire of transporters, solute carriers, channels and long non-coding RNAs (lncRNAs) across these 22 clusters (**Supplementary Fig. 4-5 and Supplementary Table 1**). This revealed in total 17 non-immune and 10 immune cell types (*PTPRC+*) (Fig. 1a-d). Where available, we provide a cell ontology ID from Ontobee (http://www.ontobee.org/)^12^.

### Proximal tubule epithelial cells

The PT is the primary site of reabsorption and filtration, where 90% of ions (Na+, Pi, HCO3-, Cl-), 65% of water and 100% of amino acids and glucose are reabsorbed into the blood. Uric acid, H+ and organic acids are removed from the blood and enter the tubular compartments, while HCO3-is reabsorbed into the interstitial tissue. These functions are specified by cells possessing different water channels, solute carriers and transporters of amino acids, glucose and urea^4^. The PT is divided into three segments (S1, S2, S3) based on ultrastructure - S1 and S2 segments encompass the convoluted portion and S3 segment the straight tubule^13^.

We define cluster 1 as PT epithelial cells (CL_0002307) as they expressed PT-specific markers such as brush border membrane components *PDZK1, LRP2*, and *CUBN*, transporters *SLC26A6*, *SLC13A2, SLC4A4, SLC5A12*, and *SLC22A6*, and *ACSM2B* among numerous other markers (**Fig. 14 1c, 2a, Supplementary Fig. 4a, 5a and Supplementary Table 1**)^14^. We did not observe distinct subclustering of the PT segments suggesting a gradual transition between these epithelial cell states. To further identify differences across these segments, we assigned 1,281 cells to S1 (CL_1000838) and 821 cells to S3 (CL_1000839) segments using known S1 (*SLC5A2, SLC2A2*) and S3 (*SLC7A13, SLC3A1*) markers and identified additional differentially expressed genes^6^. Surprisingly, plasma protein encoding genes (*e.g. ALB, RBP4, GC* and *PROC*), normally thought to be liver-derived, were expressed in PT (**Supplementary Fig. 3**)^15–17^.

PT-like cluster 5, originally a sub-set of cells within the proximal/distal/PEC (PT-DT-PEC) cluster in Fig. 1a, expressed many of the PT markers but formed a distinct cluster. We identified 3 sub-clusters within: (1) cells abundant in mitochondrial genes; (2) cells with high expression of *VIM, VCAM1, KRT18, SOCS3, ATF3* and *PDZK1;* and (3) a distal nephron - PT doublet cluster (Fig. 2a-d). PT cells in general and a subset of mouse S3 PT cells have been described to be especially abundant in mitochondria^18^. The 1^st^ sub-cluster were not dying cells as the median number of genes and transcripts were higher than cluster 1, so we hypothesize that they belong to this poorly characterized cell type (**Supplementary Fig. 2f-g**)^18^. The 2^nd^ sub-cluster marked parietal epithelial cells (PECs) (CL_1000452) lining the Bowman’s capsule, which are known to specifically express *VCAM1^19^*. We also validated the expression of VIM and other identified markers in PECs through our IMC and IHC data from Human Protein Atlas (HPA) (www.proteinatlas.org) (Fig. 2b-d **and Supplementary Fig. 7**)^20^.

**Figure 2:**
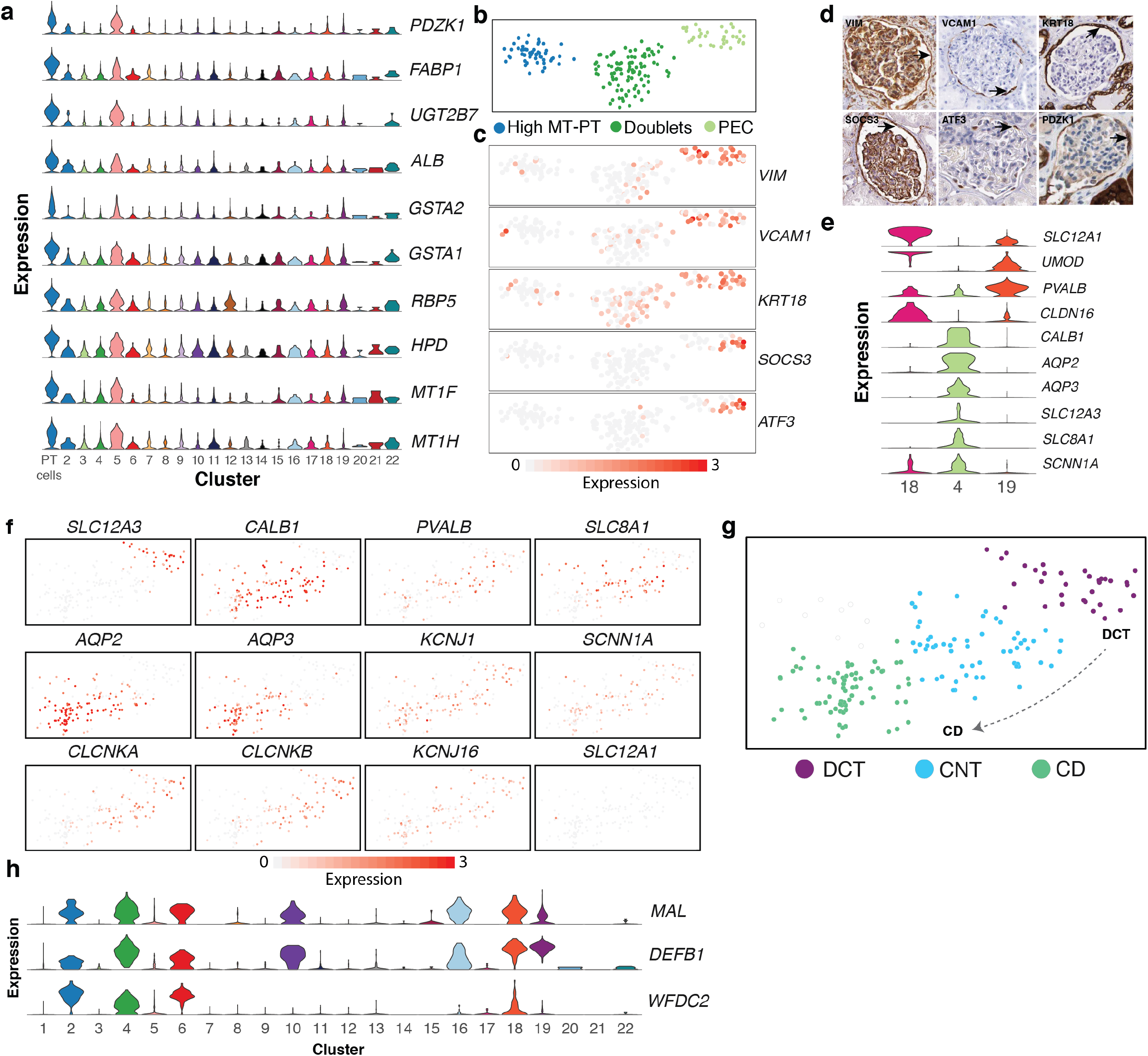
Cell types and markers of proximal tubule and distal nephron segments. (a) Violin plot of selected markers of proximal tubule epithelial cells represented by clusters 1 and 5. (b) Subclustering of cluster 5 identifies high mitochondrial PTs, PECs and doublets. (c) tSNE plots of PEC enriched markers and (d) validation IHC images from HPA (www.proteinatlas.org). Arrows indicate PECs. (e) Violin plots of markers specific to TAL, DCT, CNT and CD segments. (f) tSNE plots of cluster 4 cells showing different segment specific markers of DCT, CNT and CD. (g) Illustration of cells within cluster 4, transitioning from DCT to CNT to CD segments of the distal nephron. Expression is represented as log CPM, from 0 (grey) to 3 (orange). (h) Violin plot of *MAL, DEFB1 and WFDC2* expression across all 22 clusters. Abbreviations: DCT – distal convoluted tubule; CNT – connecting tubule; CD – collecting duct; TAL – thick ascending limb; MD – macula densa; PEC – Parietal epithelial cells; HPA – Human Protein Atlas; MT – Mitochondrial; PT – Proximal tubule epithelial cells.

### Thick ascending limb (TAL) to distal nephron (DCT to CD)

The S3 of PT transitions into the Loop of Henle (LOH), which is the main site of urine concentration where water is reabsorbed into blood. It includes the thin descending and ascending limbs and the thick ascending limb (TAL). TAL connects to the distal nephron, the site for fine regulation of various solutes and water essential in blood pressure homeostasis, and is well-delineated with known markers^5^. We identified cells of the LOH and the entire distal tubule compartment – TAL, distal convoluted tubule (DCT), connecting tubule (CNT) and collecting duct (CD) (**clusters 2, 4, 18 and 19; Fig. 1b-e, Supplementary Fig. 4 and Supplementary Table 1**).

**Cluster 2** cells expressed markers of the thin ascending limb (CL_1001016) (*ELF3, CLDN3, CLDN4, CLDN10* and *TACSTD2*), and TAL (*SLC12A1*), representing cells transitioning from the thin segment of LOH to TAL (Fig. 1c **and Sup Fig. 4b, n**)^6^. Cluster 18 cells expressed the highest levels of *UMOD* and *SLC12A1* marking them to be TAL (CL_1001106) (Fig. 2e-g)^21^. Cluster 19 represents cells transitioning from TAL to DCT (CL_1000849) as they expressed lower levels of *SLC12A1* and *UMOD* but high levels of *PVALB* (an early DCT marker/DCT1) and co-expressed *CLDN14, CLDN16* and *CLDN19* (**Fig. 1c, 2e and Supplementary Fig. 5c**)^7,22^. Since both clusters 18 and 19 express *SLC12A1*, they may contain macula densa cells (CL_1000850) within^21^.

**Cluster 4** contained a subset of cells uniquely expressing *SLC12A3*, a marker of DCT1/2 segment, followed by cells with a gradient in the expression of *PVALB, CALB1, SLC8A1, AQP2* and *AQP3*, marking the transition from DCT1/2 to CNT (CL_1000768) and CD (CL_1001431) (Fig. 2e-g)^4,21,22^. No cells co-expressed *SLC12A1* and *SLC12A3*, consistent with reports that *SLC12A1+* TAL cells abruptly transition into the DCT segment lined by *SLC12A3+* cells (Fig. 2e-g)^5^. *CLCNKB* (CIC-K2), an anion channel which mediates Cl-reabsorption, was present in TAL, DCT, CNT and intercalated cells but not in the *AQP2+, AQP3+* CD principal cells, further confirming the identity of these sub-clusters (**Fig. 2f and Supplementary Fig. 5k**)^23^. Other known apical and basolateral ion channels and transporters such as *CLCNKA, SCNN1A, KCNJ1, KCNJ16* and Na+/K+ ATPase subunits *ATP1A1, ATP1B1*, presented additional heterogeneity within the DCT, CNT and CD cells (**Fig. 2f**)^21^.

Notably, *MAL, DEFB1* and *WFDC2* mark cells of TAL, DCT, CNT and CD, with the exception of *WFDC2*, which was absent in Type A intercalated cells (Fig. 2h). Although *MAL* and *DEFB1* have been described in the context of the kidney, and *WFDC2* has been recently proposed as a serum biomarker of lupus nephritis, their cell type specific expression in the human kidney has not been well delineated^24,25^. From our data, we identify these as suitable pan-distal nephron markers.

### Intercalated cells

Intercalated cells (ICs) are specialized epithelial cells which regulate acid-base homeostasis through the exchange of various ions (Na+, Cl-, K+, HCO3- and NH3+) via ion channels and pumps. Found interspersed in the distal tubule (DCT to CD), they have paracrine signaling roles with principal cells. Three different IC types are known – Type A (IC-A), Type B (IC-B) and Non-A/Non-B ICs. IC-As are acid secreting (urinary acidification), which possess the H+ATPase vacuolar type proton pumps on the apical pole, the AE1 chloride-bicarbonate exchanger (*SLC4A1*) on the basolateral pole and lack pendrin (*SLC26A4*), a chloride-bicarbonate exchanger. IC-Bs are basic/bicarbonate-secreting cells, which express pendrin on the apical pole and H+ATPase pumps on the basolateral pole. Non-A/Non-B ICs possess both pendrin and H+ATPase pumps on the apical pole and are hypothesized to represent the intermediate/transitional state when IC-Bs switch to IC-A cell types and are located in the CNT. Dysfunction of ICs is associated with acidosis and alkalosis^24^.

We found three clusters of ICs (CL_1001432) (**Cluster 6, 10 and 16**) marked by pan-IC markers - *ATP6V0D2, ATP6V1B1* and *ATP6V1G3^24^. FOXI1*, a transcription factor specific to and required for ICs, was also detected in these populations further confirming the identity of these cells (Fig. 1b-c, 3a-b)^26^.

**Figure 3:**
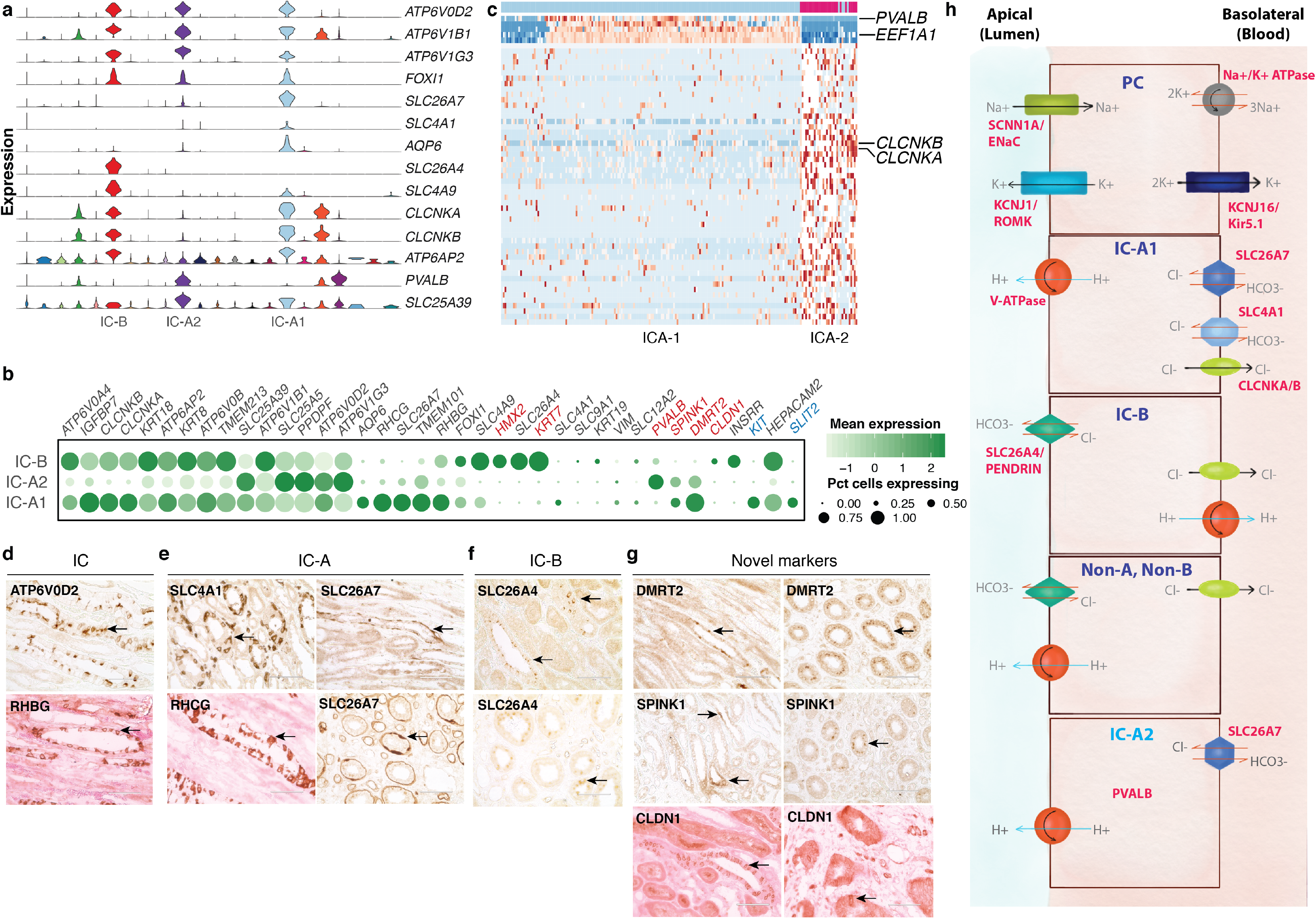
Intercalated cell types and markers. (a) Violin plot showing pan-intercalated, IC-B, IC-A1 and novel IC-A (IC-A2) subtype specific markers across all 22 clusters. (b) Bubble plot of selected IC-subtype markers. Size of dots represents percentage of cells expressing a particular maker and intensity of color represents level of mean expression. Novel markers are shown in red and signaling molecules in blue. Legends are shown below. (c) Heatmap showing scaled expression (log CPM) of discriminative marker gene sets across IC-A1 and IC-A2 (novel) subtypes, with cells as columns and genes as rows. *PVALB* and *EEF1A* are unique to IC-A2 while *CLCNKA* and *CLCNKB* are present in IC-A1 subtype. Color scheme represents Z-score distribution from −2 (blue) to 2 (dark red). (d-g) IHC on normal kidney FFPE section showing (d) intercalated cell markers, (e) IC-A markers, (f) IC-B markers, and (g) novel markers of IC-A (DMRT2 and SPINK1) and IC-B (CLDN1) subtypes. Arrows indicate respective protein expression in ICs. SPINK1 is a secreted protein, hence appears diffuse. Some sections, counter-stained with eosin, appear pink. (h) Illustration of solute carriers, channels and ATPase pumps present in PCs, IC-A1, IC-B, non-A/non-B and IC-A2 (novel) cell types. Not drawn to scale. Abbreviations: IC-B – intercalated cell type B; IC-A1 – intercalated cell type A; IC-A2 – novel intercalated cell type A; IC – intercalated cell; PC – principal cell; IHC – immunohistochemistry.

Two of these **clusters - 10 and 16** - were marked by *SLC26A7*, a marker of IC-A (CL_0005011)^27^. Genes encoding transporters/pumps revealed further heterogeneity within these IC-A subtypes (Fig 3a-c)^24^. **Cluster 16** cells (henceforth called IC-A1) co-expressed *ATP6AP2, RHCG, CLCNKA/B, SLC4A1, KIT* and high *AQP6*, which were low or absent in **cluster 10** (Fig 3a-b)^28^. **Cluster 10** cells (henceforth called IC-A2) however uniquely expressed *PVALB*, a DCT1 marker^22^. IC-A2 cells did not express *SLC26A4*, hence unlikely to be Non-A/Non-B ICs (Fig 3a-c). We denote this IC-A2 population as a novel subtype of IC-A. IC-A1 and IC-Bs had the greatest UMI counts of any cell type in our data set with median counts of 7,500 and 7,000, respectively. This is a measure of RNA content and likely reflective of cell size. ICA-2 cells were lower in median UMI detected (~4,900), still significantly higher than most all other clusters in the data set but further distinguishing these cells from the two other IC clusters, and perhaps reflecting a relatively smaller size (**Supplementary Fig. 2g**).

**Cluster 6** expressed IC-B (CL_0002201) markers *SLC26A4* and *SLC4A9* (AE4)^24^. *INSRR* was specifically expressed in IC-B and not in either IC-A subsets^29^. Similar to the IC-As, they also expressed *ATP6AP2, CLCNKA, CLCNKB*, and *RHBG* (but not RHCG)^24^. IC-B uniquely expressed high levels of epithelial marker *KRT7* (Fig. 3a-b, d and f). Apart from differences in the intracellular localization of *SLC26A4*, there are no known markers that distinguish Non-A/Non-B ICs from IC-Bs, thus both may be present within this cluster^24^.

We identified novel markers of IC-A and IC-B. *TMEM101*, an undefined transmembrane protein, *SPINK1*, a trypsin inhibitor, *DMRT2*, a transcription factor, and *C12orf75*, an undefined transcript, were all specifically expressed in IC-A1 and IC-A2 cells. IC-B uniquely expressed *CLDN1*. We verified these new markers by IHC (Fig. 3b and g). Indeed, a recent mouse study described the expression of *Spink1* and *Dmrt2* in IC-A cells^29^. They also reported *HEPACAM2* expression in IC-A, in contrast, we found *HEPACAM2* highly expressed primarily in IC-Bs and IC-A1 and lower in IC-A2 (Fig. 3b)^29^.

### Blood vessels

Endothelial cells (ECs) form the inner lining of all blood and lymphatic vessels playing critical roles in kidney architecture and function. They are the primary filtration barrier between blood and interstitial tissues, allowing exchange of nutrients, metabolites, waste and cells. Glomerular endothelial cells (GECs), which are in close contact with the podocytes and mesangial cells form the first site of filtration. The peritubular capillaries of the descending vasa recta (DVR) and ascending vasa recta (AVR), are closely knit with the nephron to allow reabsorption and secretion of ions and water, assisting in establishing counter-current exchange and concentrating the primary urine^8^. The DVR is known to be continuous and non-fenestrated unlike the fenestrated GECs and AVR^8^. Water exchange happens between the DVR-interstitium-AVR regions, hence the DVR peritubular capillaries are marked by the water channel *AQP1*. AVR and GECs can be further distinguished by *PLVAP*, a fenestrae diaphragm marker, known to be present only on AVR and not GECs or larger blood vessels^8^.

We identified three distinct clusters of ECs, marked by pan-endothelial markers *ENG* and *EMCN* (**clusters 12, 13 and 14 in Fig. 1b-c**). Other little-known EC-specific markers include *SLCO2A1*, a prostaglandin transporter, and *FXYD6*, a phosphohippolin that potentially regulates the Na+/K+ ATPase (**Supplementary Fig. 5h and j**)^30,31^. We define **cluster 12** as AVR (CL_1001131) as it expressed *PLVAP* but was low in *AQP1*, and **cluster 13** as non-fenestrated DVR (CL_1001285) as it was *PLVAP-*, but positive for *AQP1* and urea transporter *SLC14A1* (UT-B1) (Fig. 1b-c and 4b)^8^. We also noted other differences between AVR and DVR, such as mutually exclusive expression of *DNASE1L3* and *IL13RA1* in AVR, and *SOST* and *IL13RA2* in DVR. Moreover, *GSN* was higher in AVR (4-fold) while *TGFBR2* was higher in DVR (5-fold) (**Fig. 1c, 4b and Supplementary Table 1**).

**Figure 4:**
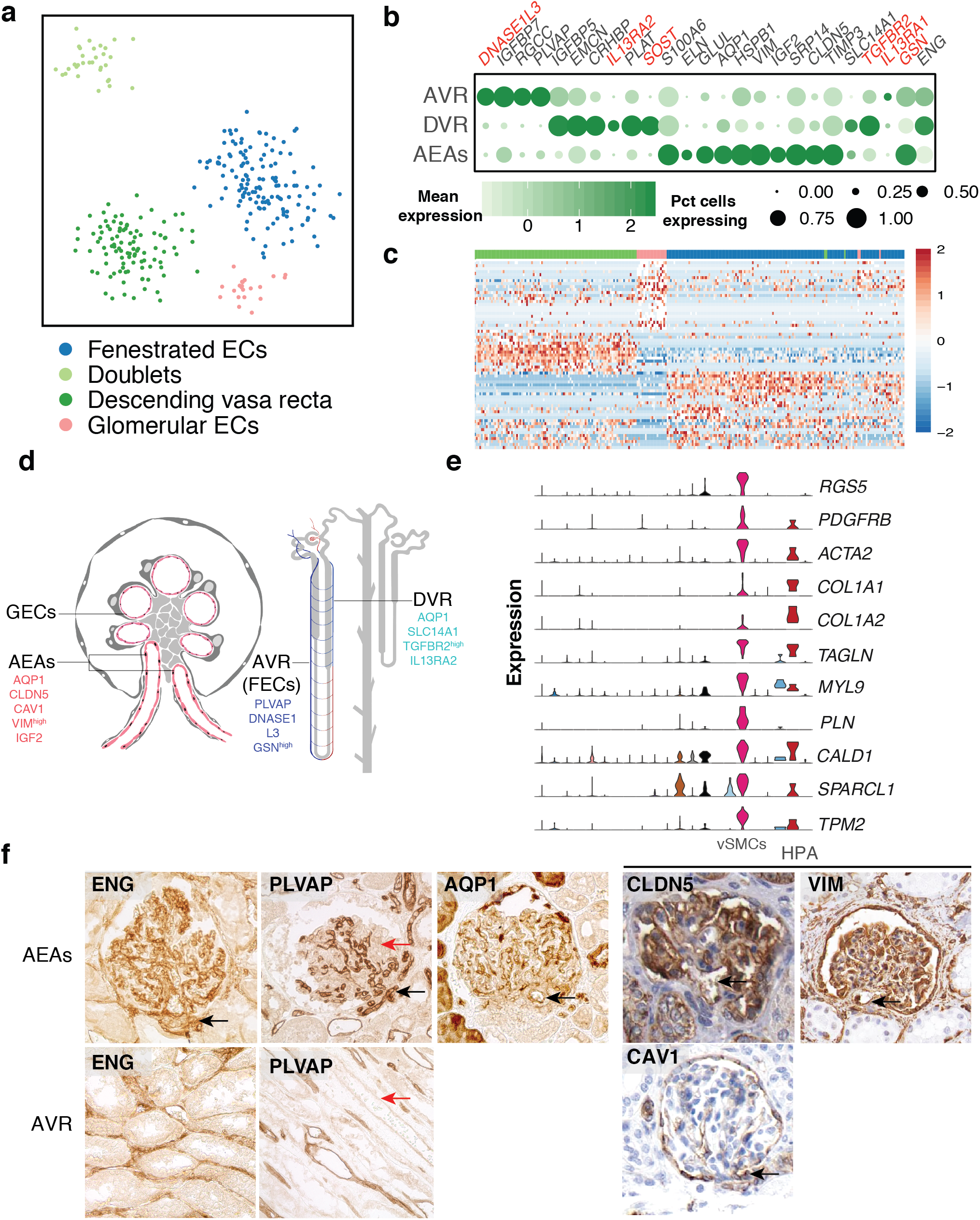
Blood vessels and vSMCs. (a) tSNE plot of AVR (FECs), DVR and AEA endothelial subtypes. A cluster of cell doublets is also shown. (b) Bubble plot of selected markers of AVR, DVR and AEAs. Size of dots represents percentage of cells expressing a particular maker and intensity of color represents level of mean expression. Disease associated markers are shown in red. Legends are shown below. (c) Heatmap showing scaled expression (log CPM) of discriminative marker genes of AVR, DVR and AEA endothelial subtypes. Cells are arranged as columns, rows are genes. Color scheme represents Z-score distribution from −2 (blue) to 2 (dark red). (d) Illustration of glomerulus with afferent and efferent arterioles and localization of AVR and DVR with respect to nephron. Not drawn to scale. (e) Violin plots showing expression of vSMC/pericyte marker genes. (f) IHC of pan-endothelial (ENG), fenestrated (PLVAP) and AQP1 expression in ECs of normal kidney section. Panel on the right shows images from HPA (www.proteinatlas.org) showing expression of CLDN5, VIM and CAV1 in ECs within glomerulus. Black arrows point to afferent/ efferent arterioles. Red arrow indicates blood vessel not marked by PLVAP. Abbreviations: ECs-endothelial cells; AVR – ascending vasa recta; DVR-descending vasa recta; AEAs – afferent and efferent arterioles; FECs – fenestrated endothelial cells; HPA – Human Protein Atlas; IHC – immunohistochemistry; GECs – Glomerular endothelial cells; vSMCs – vascular smooth muscle cells.

We define the third cluster of ECs as arteries/afferent-efferent arterioles (AEAs) (CL_1001006/ CL_1001009/ CL_1000891), based on the expression of *CLDN5, AQP1, CAV1, VIM* and low in *PLVAP. CLDN5+* ECs are known to be expressed in AEAs, arteries and podocytes but not in the kidney veins and glomerular capillaries, which we confirmed by IHC from HPA (Fig. 4b and f)^20,32,33^. *ELN* (elastin), a component of large artery extracellular matrix, *GLUL* and *IGF2* are also unique to this cluster (Fig. 4b). Notably, while publications report the lack of *PLVAP* in fenestrated GECs, our IHC analysis clearly shows *PLVAP* expression in a subset of GECs (Fig. 4b and f)^8^.

The mural cells of blood vessels - vascular smooth muscle cells (vSMCs) and pericytes, surround the vessel walls and exchange signaling cues with ECs to regulate endothelial biology^34^. **Cluster 17** consisted of cells negative for endothelial markers but co-expressing *RGS5, PDGFRB, ACTA2* and *TAGLN* – markers of pericytes (CL_1001318) and vSMCs (CL_0000359) (**Fig. 1b-c, 4e, Supplementary Fig. 4 and Supplementary Table 1**)^34^ There has yet to be definitive markers that distinguish the two and we anticipate both of these mural cells to be present – vSMCs associated with large and pericytes with small vessels – based on our IMC analysis (**Supplementary Fig. 7**).

### Glomeruli: Podocytes and mesangial cells

Podocytes are specialized epithelial cells lining the glomerular capillaries, separated from each other by a glomerular basement membrane. They possess a unique structure consisting of interdigitated podocyte foot processes with slit diaphragms, which wrap around the glomerular tuft for filtration^3^. They can be identified by podocyte-specific markers, *WT1, PODXL, NPHS1* and *SYNPO* among others^3^.

Despite the challenges associated with dissociating glomeruli and isolating podocytes (CL_1000451), we identified four podocytes which co-expressed a set of markers, including well-known podocyte markers, highly specific to these cells (**cluster 20;** Fig. 1b-c, 5a-b **and Supplementary Table 1**). Novel markers enriched in these cells, some confirmed as glomeruli localized from HPA, included *PCOLCE2, FGF1, ZDHHC6*, and *ENPEP* (Fig. 5f)^20^. Many of these markers were also found in two cells from our original version 1 analysis (**Supplementary Fig. 8**). The biological reproducibility of multiple markers across multiple cells gives us confidence in assigning these six cells as podocytes.

**Figure 5:**
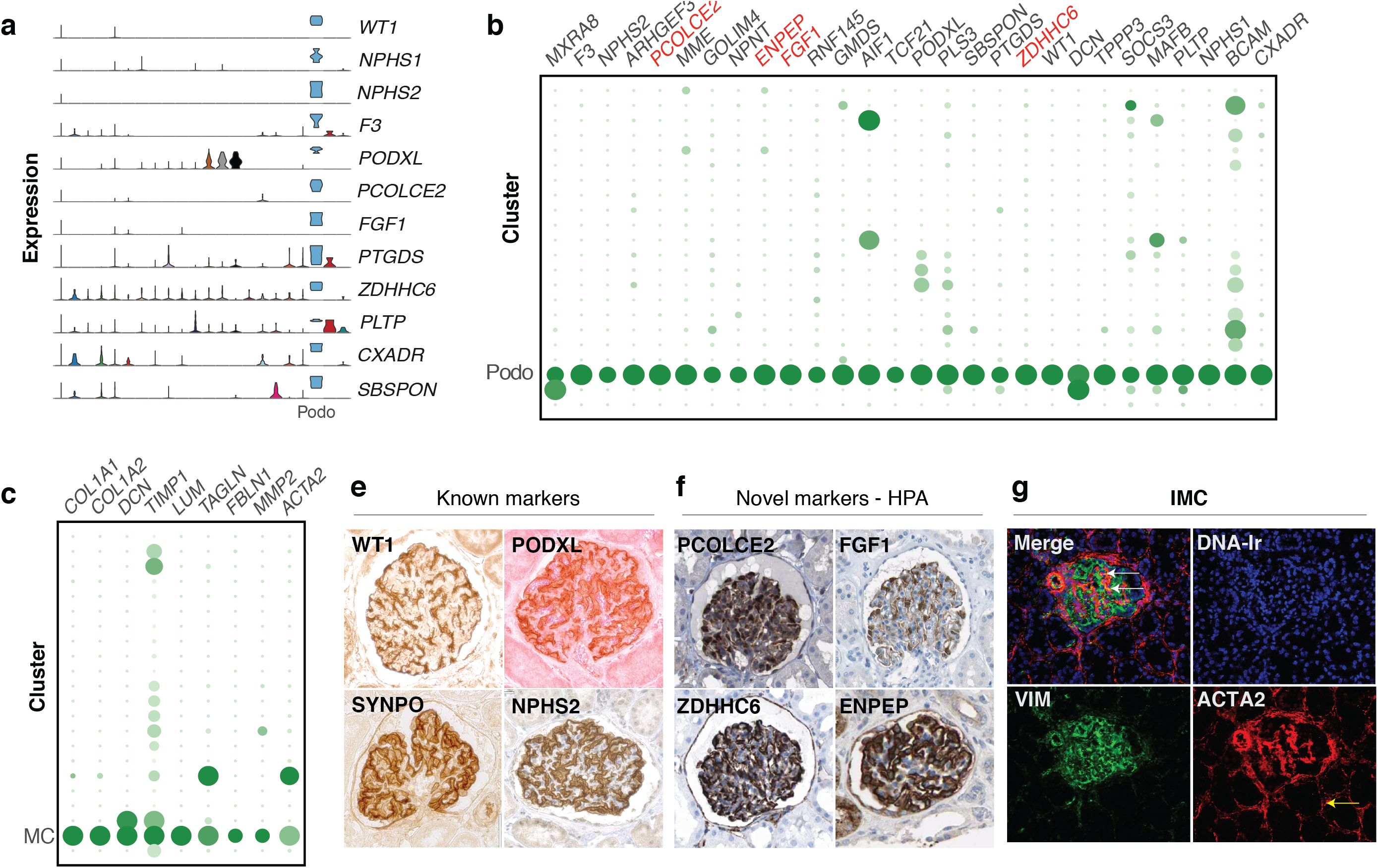
Glomerular cell types, markers and spatial localization of mesangial cells. (a) Violin plots showing the expression of podocyte marker genes. (b) Bubble plot showing genes enriched in podocytes. Novel markers of podocytes are shown in red. (c) Bubble plot showing showing genes enriched in mesangial cells. (e) IHC images validating protein expression of podocyte markers WT1, PODXL, SYNPO and NPHS2 on human kidney FFPE tissue. (f) IHC images from HPA (www.proteinatlas.org) validating protein expression of newly identified podocyte markers PCOLCE2, FGF1, ZDHHC6 and ENPEP. (g) IMC images showing co-localization of VIM and ACTA2 expression in the glomerulus (white arrow) and ACTA2 expressing pericytes (yellow arrow). Abbreviations: MC-mesangial cells; Podo-podocytes; HPA-Human Protein Atlas; IHC-immunohistochemistry; IMC-imaging mass cytomtery.

Mesangial cells are modified smooth muscle-like cells present in the glomerulus to provide structural support, basement membrane remodeling and maintenance of vascular integrity. They are highly similar in function to vSMCs/pericytes with contractile ability. Therefore, they share markers with these cell types and other known markers of fibroblasts, making it challenging to identify these cells by a combination of markers alone without spatial information^35^. They are however known to express a basal level of chemokines which are upregulated under certain conditions^36^.

We found three cells that co-expressed *COL1A1, COL1A2, TPM2, MYL9, VIM, ACTA2*, and chemokine *CCL2*, but devoid of *RGS5* (**cluster 21; Fig. 1b-c, 4e, 5c and Supplementary Fig. 5k and 7a**). Owing to the lack of known specific markers to distinguish these cells from fibroblasts, we used IMC to assess the abundance and localization of *ACTA2* and *VIM* double-positive cells. While ECs within the glomeruli are VIM-positive and ACTA2-negative, we also identified a few cells within the glomeruli that are *ACTA2/VIM* double-positive. Though many cells were positive for either of each across the entire tissue section, there were no such double-positive cells outside of the glomeruli (**Fig. 5g and Supplementary Fig. 7**). Thus we would suggest that these three cells, co-expressing many genes specific to only these cells, are likely mesangial cells (CL_0000650).

### Immune cells

The kidney is a highly vascularized tissue so we anticipated capturing immune cells of the peripheral blood. In addition, interstitial myeloid cells within the kidney - macrophages and dendritic cells - have been described^37,38^. Lymphocytes (T, NK, B and plasma), all presumed to be peripheral blood derived, were well represented in our data (65.6% of immune cells) and identified by classic markers of each cell type (**clusters 7-9 and 22,** Fig. 1b-c and 6)^39^. A cluster of 20 mast cells was also identified (**cluster 15**, Fig. 6) (CL_0000097). Two clusters of myeloid cells representing 32.9% of the immune cell population were identifiable based on the expression of CD68 (**cluster 3 and 11,** Fig. 1b and 6e)^39^.

**Figure 6:**
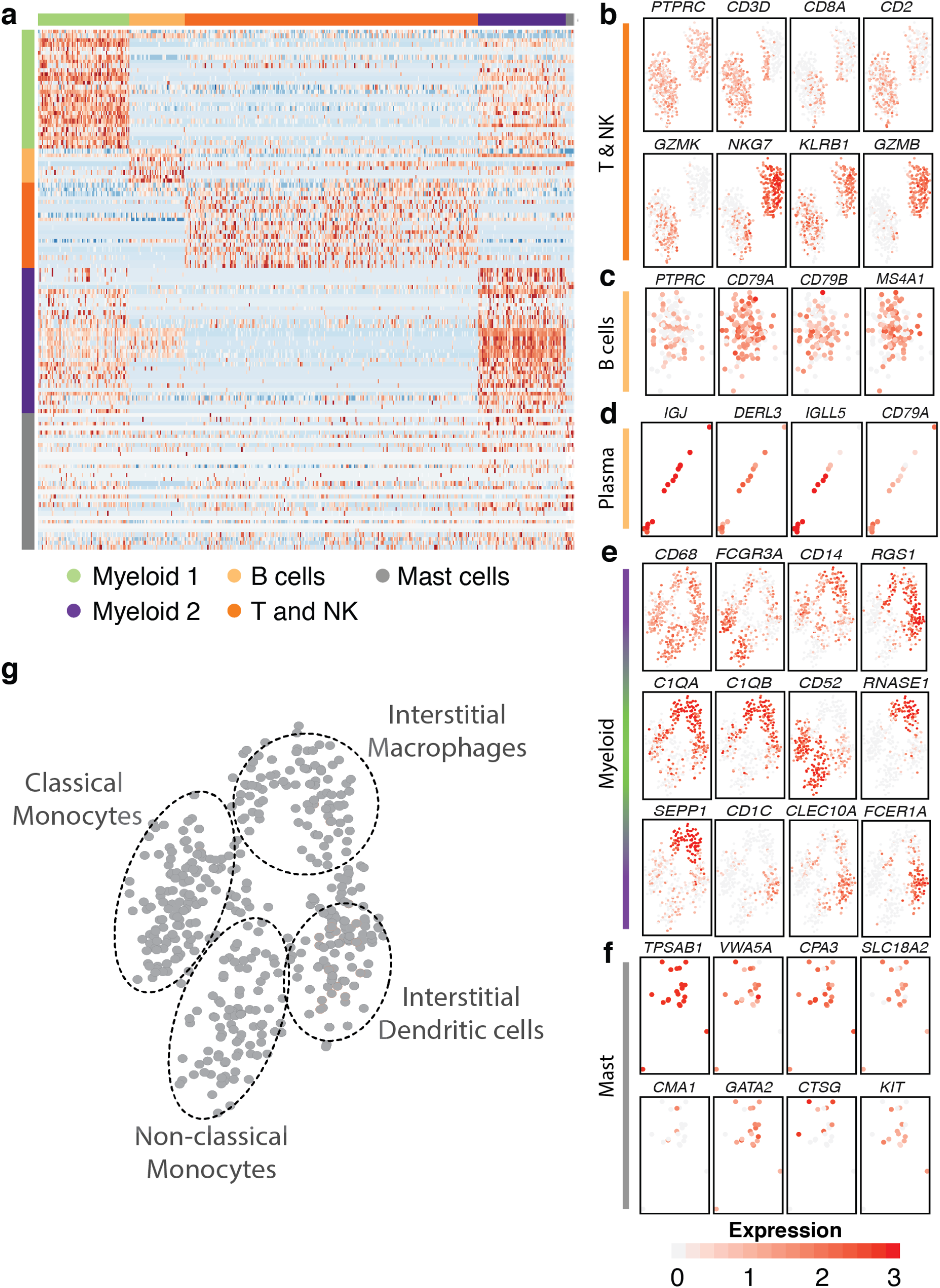
Immune cell types and markers in the kidney. (a) Heatmap showing markers of major immune cell populations. Cells are arranged as columns, rows are genes. Color scheme represents Z-score distribution from −2 (blue) to 2 (dark red). (b-f) Panel plots showing the expression of key marker genes for each major immune cell type. Expression is represented as log CPM, from 0 (grey) to 3 (orange) (g) Illustration of different cell subsets within the combined myeloid cell cluster.

To better characterize the myeloid lineage cells, we combined the two *CD68+* clusters for re-analysis. Prior to this, it was apparent that a major contributor differentiating these two clusters was the expression of *CD52*, high in **cluster 3** (and additionally expressed in the T/NK/B/mast cell clusters) and low in **cluster 11**. CD52 is a known marker of peripheral blood cells and, as a highly negatively charged molecule, speculated to function in inhibiting adhesion. It has also been described to be absent on tissue-resident T cells and DCs^40,41^. This suggested *CD52+* **cluster 3** represents monocytes of the peripheral blood and *CD52-* **cluster 11**, the interstitial myeloid cells. Subsequent analysis showed *CD52* low/negative cells containing markers of interstitial macrophage and dendritic cells (DC) (Fig. 6e, g), specifically a sub-population marked by *CD1C, CLEC10A* and *FCER1A*^39^ which we define as *CD1c* DCs (CL_0002399), and another subset expressing high levels of the tissue-resident macrophage marker *C1QA^42^. RNASE1* and *SEPP1* also marked this *C1QA+* population, and *RGS1* more broadly the *CD52* low/negative cells. Thus, within the myeloid compartment we define clusters of classical (CL_0000860) and non-classical monocytes (CL_0002396) in addition to interstitial macrophage and DCs (Fig. 6e, g) and provide evidence for the utility of *CD52* in differentiating between peripheral blood and interstitial immune cells.

### Receptor-ligand interactions

We explored potential communication axes between cell types by investigating the expression patterns of ligands and receptors in all the cell types identified in our data (Fig. 7a-b). While numerous ligand-receptor interactions could be discerned from our analysis, immune cells (myeloid lineage and T/NK/B lymphocytes) were the largest receivers of ligands, followed by ECs and intercalated (IC-A, IC-B) cells. On the contrary, *PVALB+* DCT1 segment and ICA-2 (novel intercalated cells) were predominantly ligand presenters (Fig. 7c). Of particular interest are interactions between: (1) vSMC/pericyte and endothelial cells (*PDGFRB-PDGFB)^43^;* (2) DCT2-CNT-CD segments and ICs (KITLG-KIT)^29^; (3) podocytes and ECs (*VEGFA - FLT1/KDR)^44^;* and (4) IC-As and ECs (*SLIT2-ROBO4*) (Fig. 7a-h)^45–47^. Our analysis identified possible signaling axes that may be important for proper kidney function and potentially disrupted in kidney pathologies.

**Figure 7:**
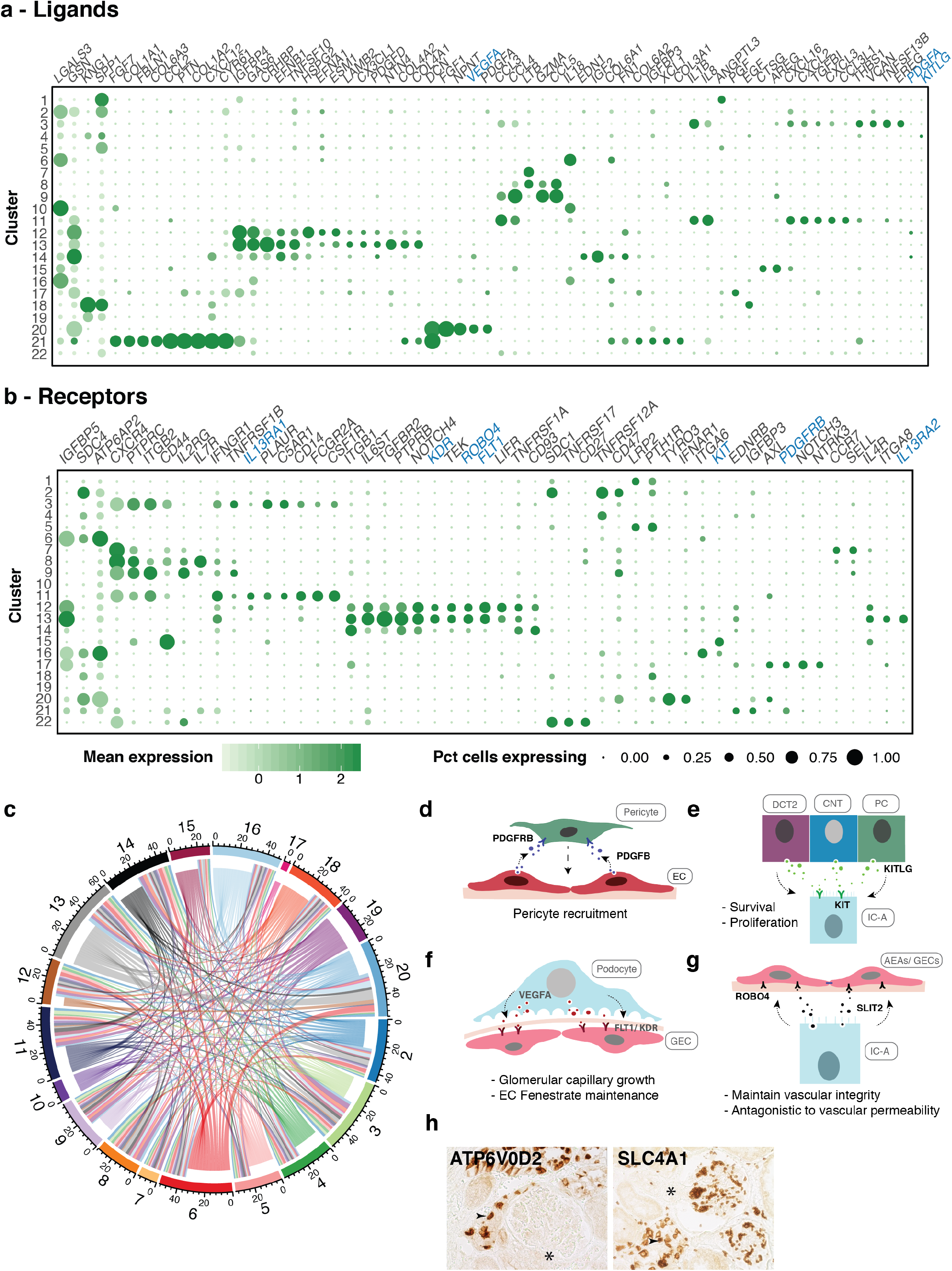
Receptor-ligand interactions in the kidney. (a) Bubble plots showing the mean expression level of selected ligand and (b) receptor encoding genes across all clusters. Signaling molecules discussed in text are shown in blue. Size of dots represents percentage of cells expressing a particular maker and intensity of color indicates level of mean expression. Legends are shown below. (c) Circos plot depicts potential interactions between ligand and receptor pairs found in our dataset. Inner ring represents broadcasting ligands, outer ring represents presenting receptors. Numbers correspond to cluster numbers. Illustrations depicting potential signaling interactions between (d) pericytes and endothelial cells, (e) IC-A and DCT2, CNT and PCs, (f) podocytes and glomerular endothelial cells, (g) IC-A and glomerular/afferent-efferent arteriole endothelial cells. (h) IHC of pan-IC and IC-A markers showing presence of ICs in close proximity to AEAs. * indicates AEAs. Abbreviations: EC-endothelial cell; IC-A-intercalated cell type A; DCT2-distal convoluted tubule 2; CNT-connecting tubule; PC-principal cell; GEC-glomerular endothelial cell; AEAs-afferent-efferent afterioles; IHC-immuno histochemistry

### Long non-coding RNAs

Long non-coding RNAs (lncRNA) display cell-type-specific expression and thus have utility in distinguishing cell types^48^. Our analysis uncovered cell-type specific expression of lncRNAs in ECs (*AC011526.1/PCAT19*), podocytes (*LINC00839, RP11-550H2.2*) and B cells (*LINC00926)^49^*. ICs expressed *CTB27N1.1/LINC01187* (IC-A, IC-B) and *PART1* (IC-A1/2), whereby *PART1* has been shown to be upregulated in chromophobe renal cell carcinoma which originates from ICs^50^.

Intriguingly, the widely studied *MALAT1* and *NEAT1* were abundantly expressed in all clusters except distal tubule, PT and ICA-2 (**Supplementary Fig. 6**). Their distinct expression, or lack thereof, in specific cell-types argues for further investigation into their functions in the kidney.

### Kidney disease genetic risk genes associated with distinct cell types

Numerous studies have identified causal genes and their variants associated with various kidney diseases^10,11^. We examined the expression patterns of causal genes involved in chronic kidney disease (CKD and estimated by serum creatinine, eGFRcrea), albuminuria, IgA nephropathy, nephrolithiasis and lupus nehpritis. Kidney disease-associated genes from GWAS were largely localized in proximal tubule, podocytes, endothelial and myeloid cells. Our analysis confirmed the cell type specific expression of CKD causal gene *UMOD* in TAL/DCT^51^, Idiopathic Membranous Nephropathy causal gene *PLA2R1* in podocytes^52^ and eGFRcrea causal gene *DACH1* in podocytes^53^. It also revealed the cell type specific expression of numerous other kidney disease associated genes which were previously unknown - *NAT8* (CKD) in PT cells, *IGFBP5* (eGFRcrea) in IC-A and ECs, *CUBN* (albuminuria) in PT cells, and *NOTCH4* and *DNASE1IL3* (lupus nephritis) in ECs (Fig. 8)^10,11,54^.

**Figure 8:**
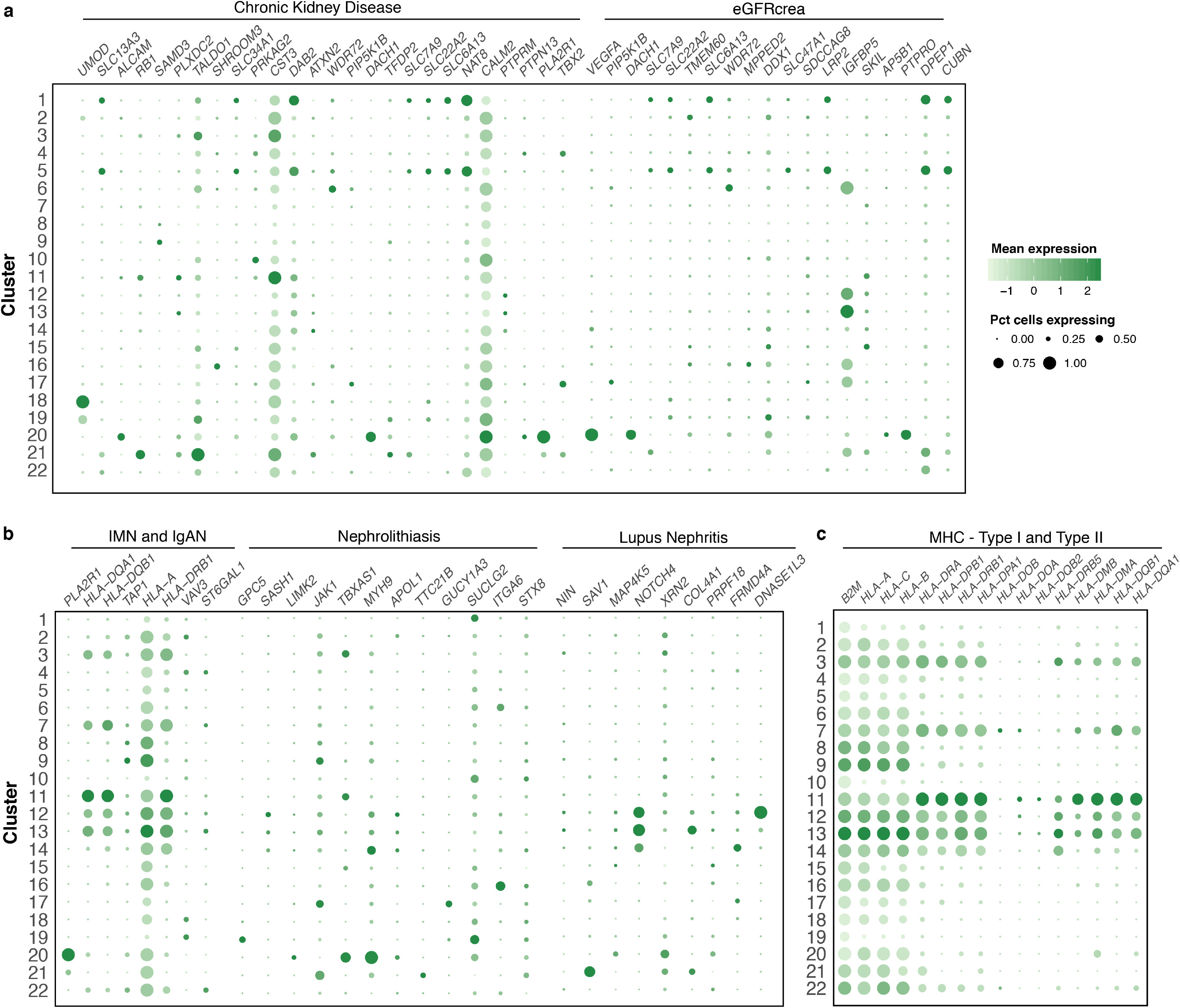
Cell type specific expression of causal genes in kidney pathologies. (a and b) Bubble plot showing the mean expression of genes associated with different kidney diseases identified in genome wide association studies (GWAS). (c) Bubble plot showing the mean expression of MHC type I and type II genes. Size of dots represents percentage of cells expressing a particular maker and intensity of color indicates level of mean expression. Legends are shown in panel (a). Abbreviations: eGFRcrea-estimated glomerular filtration rate creatinine; IMN-Idiopathic membranous nephropathy; IgAN-IgA nephropathy; MHC – Major histocompatibility complex.

An interesting side note on HLA molecules, included in this analysis, identified HLA class II molecules, generally abundant in professional antigen presenting cells, expressed on all ECs. In addition, MHC Class I molecules (*HLA-A, HLA-B, HLA-C, B2M*), which are believed to be expressed in all nucleated human cells, were strikingly low in PT, DCT1 and IC-A2 (Fig. 8c).

## Discussion

We present an unbiased single cell analysis of the human kidney to define cell-type specific transcriptomes. Previous single cell transcriptome studies on kidney were mostly performed in mice and/or were focused on a specific kidney cell type (mesangial, podocytes, principal and intercalated cells)^29,55–57^. Our unbiased analysis has identified 27 different cell types including 17 non-immune and 10 immune cell types. We have confidently mapped the transcriptome of the nephron, collecting duct cell types and interstitial macrophages and DCs based on uniquely expressed markers. In addition, we describe cell type-specific expression of solute carriers, channels, transporters, receptors and ligands that will be useful to delineate the multitude of functions performed by the various kidney cells. We also identified new markers and lncRNAs uniquely expressed in specific cell types such as IC subtypes, the distal nephron, podocytes and endothelial cells.

### Novel cell populations in the human kidney

We discerned the heterogeneity within various cell types, such as the intercalated (IC-A1, IC-A2 and IC-B) and endothelial (AVR, DVR and AEAs) cells. We identified a novel IC-A subtype (IC-A2) and a *CLDN5+* endothelial population. The novel ICA-2 expresses *PVALB*, which functions to buffer Ca2+ and Mg2+ ions in the distal nephron. We postulate that since *PVALB* is well-known to mark the early DCT, the IC-A2 subtype likely resides in this segment. Although the IC-A2 subset was present in two different kidney samples, whether this cell type represents a transitional state present only under certain physiological conditions remains to be verified. *CLDN5*+ ECs have been described before in kidney, we show here that they segregate as a separate subtype of ECs. *CLDN5* is a tight junction protein expressed in microvascular ECs forming endothelial barriers of lung, brain (blood-brain-barrier), retina, spinal cord and kidney, and decrease in its expression is associated with increase in paracellular permeability^58–60^. Although GECs are known to be fenestrated but lack *PLVAP*, we found *PLVAP* expression in a subset of these ECs along with *CAV1* and *AQP1*, indicating that GECs use multiple modes of transcellular transport^61^.

### Protective and survival signaling between the kidney cell types

In addition to cell type classification, we also investigated cell-to-cell communications. While PDGFB-mediated recruitment of pericytes to ECs is well known, KIT-KITLG signaling in ICs has only been recently described^29,43^. Chen et al (2017) described the expression of KIT receptor in IC-A, and IC-B (weakly) cells and its ligand, KITLG, in principal cells^29^. We detected KIT in IC-A but none in IC-B or novel IC-A2 cells, and KITLG was expressed in PCs as well as DCT2 (*SLC12A3+*) and CNT in human kidney. KITLG-KIT tyrosine kinase receptor signaling activates the MAPK, PI3K, JAK-STAT and PLC-PKC pathways. KIT signaling is therefore associated with cell proliferation, survival, adhesion and chemotaxis^62,63^. We hypothesize that the KITLG secreted by cells of these distal nephron, where IC-A are found interspersed, plays a role in their survival and proliferation.

Another notable interaction was among podocytes, ICs and ECs. VEGFA-FLT1/KDR signaling axis between podocyte and ECs is essential for glomerular capillary growth and maintenance of endothelial fenestrae^44^. On the other hand, SLIT2-ROBO4 function antagonistic to VEGF-mediated vascular permeability, whereby they upregulate *CLDN5* and *CDH5* in ECs, thus impeding paracellular transport and maintaining vascular integrity^45^. Intriguingly, IC-A cells expressed *SLIT2*, which encodes a secreted glycoprotein. We postulate that ECs in reasonable proximity to IC-As, such as AEAs will be subject to this restriction to vascular permeability. Indeed, our histology analysis shows the presence of ICs close to AEAs, indicating SLIT2-ROBO4 signaling may be responsible for *CLDN5* induction in these ECs. Moreover, several studies have implicated a role for SLIT2-ROBO4 signaling in the regulation of vascular and renal injury mediated through infiltrating immune cells^46,47^. While ICs are known for their role in immune response against bacterial infection, *SLIT2* expression by IC-A implicates these cells in a potentially new and alternative protective role in kidney injury^24^.

### Unexpected expression of plasma protein encoding genes in proximal tubule cells

Interestingly, we found that PT cells expressed plasma protein encoding genes (*ALB, RBP4, GC*, and *PROC*), which are reported to be synthesized and secreted by the liver^15–17^. Urinary ALB and RBP4 are biomarkers of albuminuria and CKD (nephritis) and renal Fanconi syndrome, respectively^10,17^. The presence of albumin in the urine is an indication of glomerular and tubular injury owing to dysfunction in filtration. Likewise, elevated serum and urinary RBP4 levels are attributed to increased RBP4 synthesis in the liver as well as a failure in proper reabsorption of RBP4-retinol complex by PT cells^64^. Our observation of *ALB* and *RBP4* expression in the PT cells raises important questions regarding the source of urinary ALB and RBP4, and perhaps indicates issues not only with reabsorption but also cellular damage of PT cells in kidney diseases. Expression of these plasma protein genes in the kidney warrants further investigation into its potential role as an alternative site for plasma protein production.

### Dysfunction of endothelial cells in kidney pathologies

Our analyses suggest an underappreciated role for ECs in a wide range of kidney pathologies. Several of the genes specifically expressed in the ECs (*i.e. GSN, DNASE1L3, IL13RA1, IL13RA2, TGFBR2*) are associated with various kidney diseases. For instance, unique mutations within *GSN* lead to a kidney-specific form of gelsolin amyloidosis. A loss-of-function (LOF) variant in *DNASE1L3* is a cause of a familial form of systemic lupus erythematosus (SLE) correlating with a high frequency of lupus nephritis^54^. Other LOF mutations in this gene have been associated with hypocomplementemic urticarial vasculitis syndrome (HUVS), which is partly characterized by glomerulonephritis^65^. *IL13RA1* and *IL13RA2* have been implicated in tissue fibrosis. *IL13RA2* has been implicated in the induction of TGFB1-mediated fibrosis in colitis and pulmonary fibrosis^66,67^. The ligand, TGF-beta is known to strongly promote renal tubulointerstitial fibrosis. Interestingly, conditional ablation of *TGFBR2* in mouse kidney interstitial cells, the cell type considered to be mediating fibrosis, did not reduce fibrosis in a kidney injury model^68^ suggesting an alternative cell type is involved. We hypothesize that TGF-beta induced fibrosis in the kidney intimately involves this *TGFBR2^high^* and *IL13RA2+* DVR endothelial cell population. Thus, endothelium restricted expression for these genes implies a central role for the peritubular capillaries in the associated pathologies.

In summary, the detailed analyses of the kidney cell types that we present in this study will serve as an important resource that will allow for a comprehensive understanding of kidney function and specific dysfunction in kidney pathologies.

## Materials and Methods

### Procurement of kidney biopsies and FFPE sections

Freshly resected renal biopsies and corresponding FFPE sections were obtained from UConn Health Research Biorepository from de-identified consented patients (male and female, age – 62-66) undergoing kidney nephrectomy for clear cell renal cell carcinoma (ccRCC). These samples were determined not to be human subjects research by the JAX IRB (IRB Reference Number: 2017-019). Only biopsies from “normal” regions (assessed by pathologists) of the kidney were utilized in this study.

### Tissue dissociation, Ab staining and FACS

Kidney tissue was processed by enzymatic dissociation with Collagenase IV (600U/mL) and DNAse (2ug/mL) for a maximum of 45min and neutralized in collection buffer (1% BSA, 10%FBS, 2mM EDTA in RPMI). Cells were spun down and treated with ACK lysis buffer (A1049201, GIBCO) for 3 min on ice and washed with collection buffer. Cells were then stained for Calcein AM (C3100MP) and DAPI (Lifetech D1306, 1ug/mL) and sorted by Fluoresence-Activated Cell sorting (FACS) to select only live cells for downstream single cell transcriptomic analyses.

### Single cell RNA-sequencing

Sorted cells were washed once with 0.04% BSA in PBS and counted on Countess II automated cell counter (ThermoFisher). 12,200 cells were loaded per lane on the 10x Chromium platform and processed for cDNA synthesis and library preparation as per manufacturer’s protocol. Kidney 1 sample was processed using Chromium™ version 1 chemistry while Kidney 2 and Kidney 3 samples were processed by version 2 chemistry. cDNA and libraries were checked for quality on Agilent 4200 Tapestation and quantified by KAPA qPCR before sequencing on a single lane of a HiSeq4000 (Illumina).

### scRNA-seq analysis

Cellranger v1.3 was used to convert Illumina base call files to FASTQ files. These FASTQ files were aligned to the hg19 genome and transcriptome provided by 10X genomics. The gene vs cell count matrix from Cellranger was used for downstream analysis. Cells with less than 500 transcripts and genes with less than 2 counts in 3 cells were discarded. The top 1000 genes with the most variance were identified based on their mean expression in the population and dispersion (variance / mean expression). Genes were binned into 50 different bins based on their mean expression and dispersion scaled with respect to the median dispersion in each bin. These genes were used to reduce the dimensions of the dataset using Barnes Hut t-SNE using default parameters. Cells were clustered in t-distributed stochastic neighbor embedding (tSNE) space using density-based spatial clustering of applications with noise (DBSCAN). Marker genes were identified using area under a receiver operating characteristic curve (AUROC) analysis. All genes that are greater than 2-fold expressed in the cluster compared to the rest of the population were analyzed using AUROC. Genes that had greater than 85% classification score were defined as markers specific to the cell type. Kidney 1 data analysis was performed separately since it was processed by version 1 chemistry. We identified the same set of cell types and markers in the kidney sample 1 as kidney 2 and kidney 3 (**Supplementary Fig. 8**).

### Ligand-receptor pair analysis

We used a list of receptors and ligand pairs from published databases^69^. Any ligand or receptor was classified as expressed if greater than 50% of the cells had non-zero counts within each cell type. To prevent spurious detection of ligand-receptor interactions due to extremely high or low cell numbers, we disregarded PT cells (cluster 1), podocytes (cluster 20), mesangial cells (cluster 21) and plasma cells (cluster 22). This is reflected in the interaction plot in Fig. 7c, which was rendered using the R package circlize.

### Immunohistochemistry

FFPE sections of 5 micron thickness were obtained from UCONN biorepository. Sections were prewarmed on slide warmer for 10min at 55^o^C and de-paraffinized using Histoclear (HS-200, National diagnostics). Sections were then rehydrated through an ethanol gradient (100%, 95%, 90%, 70% for 3 min each) and rinsed once in water. Antigen retrieval was then performed either in alkaline (BSB 0030, BioSb) or R-universal buffer (AP0530-500, EMS) (121^o^C, 15min) in a TintoRetriever pressure cooker (BSB 7008). Immuno-staining was then performed using BioSB kit (Mouse/Rabbit PolyDetector Plus DAB HRP Brown Detection System, BSB 0257) as per manufacturer’s instructions with a minor change - primary Ab was incubated overnight at 4^o^C. List of antibodies used are provided in Supplementary Table 2.

### Imaging Mass Cytometry

FFPE sections (5um thick) were stained with metal-tagged antibody cocktail as per the protocol described by Chang et al 2017, with some modifications^70^. Briefly, tissue sections were rehydrated as outlined above and heat-mediated antigen retrieval was performed in citrate buffer (BSB 0023) in the TintoRetriever pressure cooker (121^o^C, 7 min). Slides were then blocked for 45 mins at room temperature with PBS containing 5% FBS and 3% BSA. Following blocking, sections were incubated with a cocktail of metal-tagged antibodies (Supplementary Table 2) overnight at 4^o^C. Slides were then washed and stained with Iridium intercalator (201192B, Fluidigm) at a final concentration of 0.25uM for 30 min at RT. Slides were then washed with distilled water and air dried. The slide was then loaded on the Hyperion™ system (Fluidigm), regions of interest were identified and tissue ablation was performed. The resulting .mcd files were exported as .tiff through MCDViewer (Fluidigm).

### Data availability

The datasets generated during and/or analysed during the current study are available in the GEO and SRA repository. These are currently embargoed until publication. SRA Study Accession ID: SRP126175, SRA Run Accession ID: SRR6348583, SRR6348584, SRR6348585, SRR6348586.

### Code availability

All the code to process, analyze and generate figures will be made available on github as ipython notebooks: https://github.com/mohanbolisetty/scRNASeq-Kidney.

## Acknowledgements

We thank Vinod Yadav and Krishna Karuturi for providing a curated ligand-receptor list, Lori Perpetua from UCONN Biorepository for assistance with samples, Fluidigm for contributing antibodies for IMC and IHC, Anthony Carcio from JAX FACS core facility. This work was supported by the JAX Scientific Services Innovation Fund and by laboratory startup funds to PR. All the authors read and approved the manuscript. All the authors declare no conflicts of financial interest.

## Author contributions

VS, MB, SS and PR conceived the study, analyzed data and wrote the manuscript. VS, MB, SS, SB and DR performed the experiments. MB performed all computational analyses. Project administration and funding acquisition: PR.

## Additional information

Supplementary figures and supplementary table 1 are attached to this paper.

## Supplementary Figure Legends

**Supplementary Fig. 1: Workflow for cell type discovery and identification in the human kidney.**

Normal human kidney biopsies were dissociated into single cells prior to profiling with 10X Chromium, followed by unsupervised analysis for cell type identification. Validations of cell type specific markers were performed on kidney FFPE sections using traditional IHC and multi-parameter IMC methods. Abbreviations: IHC-immunohistochemistry; FACS-Fluorescence-activated cell sorting; IMC-imaging mass cytometry; FFPE-Formalin fixed paraffin embedded.

**Supplementary Fig. 2: Quality and clustering metrics**. (a) Histogram of genes and (b) transcripts detected in the 19,341 cells in the kidney data. (c) Technical variables like genes and (d) transcripts detected doesn’t drive the unsupervised clustering of cells. (e) Cells from kidneys 2 and 3 are present in all clusters and is not patient specific. (f) Distribution of genes and (g) transcripts detected in each cluster in kidney data.

**Supplementary Fig. 3: S1 and S3 segments of the proximal tubule**. (a) Violin plot of selected markers of S1 and S3 segments of the proximal tubule epithelial cells. (b) IHC validation of S3 specific genes from HPA (<u>www.proteinatlas.org</u>).

**Supplementary Fig. 4: Markers of the cell types in the human kidney**. (a – o) Bubble plot showing selected cell type specific markers across all 22 clusters. Size of dots represents percentage of cells expressing a particular maker and intensity of color indicates level of mean expression. Legends are shown below. Abbreviations: NK – natural killer; IC-A – intercalated cell type A; IC-B – intercalated cell type B; EC – endothelial cell; Pct – percentage; PT – proximal tubule; AVR – ascending vasa recta; DVR – descending vasa recta; FEC – fenestrated endothelial cells.

**Supplementary Fig. 5: Expression of channels and transporters in the cell types of the human kidney**. (a – l) Bubble plot showing selected cell type specific markers across all 22 clusters. Size of dots represents percentage of cells expressing a particular maker and intensity of color indicates level of mean expression. Abbreviations – MHC – Major histocompatibility complex.

**Supplementary Fig. 6: Expression of long non-coding RNAs in the cell types of the human kidney**. (a) Bubble plot showing selected cell type specific markers across all 22 clusters. Size of dots represents percentage of cells expressing a particular maker and intensity of color indicates level of mean expression.

**Supplementary Fig. 7: Multi-parameter protein profiling of human kidney section using IMC.** (a) FFPE human kidney section of 5um thickness was stained with metal-conjugated antibodies for ACTA2, VIM, EPCAM and DNA-Ir intercalator. A region of the section (l x w: 12,000 micron x 500 micron) extending from the cortex to the medullary region of the kidney was laser ablated using the Hyperion™ system (Fluidigm). (b) Magnified image of a glomerulus, boxed in (a), showing merged image of ACTA2, VIM, EpCAM and DNA-Ir. Individual markers are shown as separate panels. Mesangial cells (white arrow) and pericytes (yellow arrow) co-express ACTA2 and VIM, while endothelial cells stained for VIM only. Parietal epithelial cells stained for VIM but not EpCAM (red arrows). DNA-Ir stained all nuclei. (c) Magnified image of renal artery, boxed in (a), showing VIM expression on the innermost layer of endothelial cells (red arrow). vSMCs surrounding the artery costained for ACTA2 and VIM ( yellow arrow). Some of the tubules were marked by EpCAM. Abbreviations: FFPE – Formalin-fixed paraffin-embedded; DNA-Ir – DNA-Iridium intercalator; vSMC – vascular smooth muscle cell.

**Supplementary Fig. 8: Human kidney cell types delineated by single cell transcriptome using the Gemcode chemistry**. (a) Unsupervised clustering of human kidney represented as a tSNE plot. Different cell type clusters are color coded. (b) Unsupervised clustering of non-PT epithelial cells of the kidney. Intercalated cells and collecting duct cells from (a) were merged and re-clustered (unsupervised), revealing finer subsets. Different cell type clusters are color coded. (c) Bubble plot showing selected cell type specific markers across all clusters. Size of dots represents percentage of cells expressing a particular maker and intensity of color indicates level of mean expression. Legends are shown below. (d) Histogram of genes and (e) transcripts detected this kidney. (f) Distribution of genes and (g) transcripts detected in each cluster in kidney data. Abbreviations: IC-A – intercalated cell type A; IC-B – intercalated cell type B; PT – proximal tubule; AVR – ascending vasa recta; DVR – descending vasa recta, CD – Collecting Duct.

## References

1 Neuen, B. L., Chadban, S. J., Demaio, A. R., Johnson, D. W. & Perkovic, V. Chronic kidney disease and the global NCDs agenda. BMJ Global Health 2, doi:10.1136/bmjgh-2017-000380(2017).

2 2017 Annual Data Report Highlights, <https://www.usrds.org/adrhighlights.aspx> (2017).

3 Dressler, G. R. The cellular basis of kidney development. Annual review of cell and developmental biology 22, 509–529, doi:10.1146/annurev.cellbio.22.010305.104340 (2006).

4 Nielsen, S. et al. Aquaporins in the kidney: from molecules to medicine. Physiological reviews82, 205–244, doi:10.1152/physrev.00024.2001 (2002).

5 Biner, H. L. et al. Human cortical distal nephron: distribution of electrolyte and water transport pathways. Journal of the American Society of Nephrology : JASN 13, 836–847 (2002).

6 Lee, J. W., Chou, C. L. & Knepper, M. A. Deep Sequencing in Microdissected Renal Tubules Identifies Nephron Segment-Specific Transcriptomes. Journal of the American Society of Nephrology : JASN 26, 2669–2677, doi:10.1681/ASN.2014111067 (2015).

7 Yu, A. S. Claudins and the kidney. Journal of the American Society of Nephrology : JASN 26, 11–19, doi:10.1681/ASN.2014030284 (2015).

8 Aird, W. C. Phenotypic heterogeneity of the endothelium: II. Representative vascular beds. Circulation research 100, 174–190, doi:10.1161/01.RES.0000255690.03436.ae (2007).

9 Giesen, C. et al. Highly multiplexed imaging of tumor tissues with subcellular resolution by mass cytometry. Nature methods 11, 417–422, doi:10.1038/nmeth.2869 (2014).

10 Wuttke, M. & Kottgen, A. Insights into kidney diseases from genome-wide association studies. Nature reviews. Nephrology 12, 549–562, doi:10.1038/nrneph.2016.107 (2016).

11 Vezzoli, G., Terranegra, A., Arcidiacono, T. & Soldati, L. Genetics and calcium nephrolithiasis. Kidney international 80, 587–593, doi:10.1038/ki.2010.430 (2011).

12 Ong, E. et al. Ontobee: A linked ontology data server to support ontology term dereferencing, linkage, query and integration. Nucleic acids research 45, D347– D352, doi:10.1093/nar/gkw918 (2017).

13 Desgrange, A. & Cereghini, S. Nephron Patterning: Lessons from Xenopus, Zebrafish, and Mouse Studies. Cells 4, 483–499, doi:10.3390/cells4030483 (2015).

14 Habuka, M. et al. The kidney transcriptome and proteome defined by transcriptomics and antibody-based profiling. PloS one 9, e116125, doi:10.1371/journal.pone.0116125 (2014).

15 Essalmani, R. et al. Thrombin activation of protein C requires prior processing by a liver proprotein convertase. The Journal of biological chemistry 292, 10564–10573, doi:10.1074/jbc.M116.770040 (2017).

16 Bernardi, M., Ricci, C. S. & Zaccherini, G. Role of human albumin in the management of complications of liver cirrhosis. Journal of clinical and experimental hepatology 4, 302–311, doi:10.1016/j.jceh.2014.08.007 (2014).

17 Jing, J. et al. Chronic Kidney Disease Alters Vitamin A Homeostasis via Effects on Hepatic RBP4 Protein Expression and Metabolic Enzymes. Clinical and translational science 9, 207–215, doi:10.1111/cts.12402 (2016).

18 Forbes, M. S., Thornhill, B. A., Galarreta, C. I. & Chevalier, R. L. A population of mitochondrion-rich cells in the pars recta of mouse kidney. Cell and tissue research 363, 791–803, doi:10.1007/s0044-015-2273-x (2016).

19 Ogawa, T. et al. Ultrastructural localization of vascular cell adhesion molecule-1 in proliferative and crescentic glomerulonephritis. Virchows Archiv : an international journal of pathology 429, 283–291 (1996).

20 Uhlen, M. et al. A human protein atlas for normal and cancer tissues based on antibody proteomics. Molecular & cellular proteomics : MCP 4, 1920–1932, doi:10.1074/mcp.M500279-MCP200 (2005).

21 Mount, D. B. Thick ascending limb of the loop of Henle. Clinical journal of the American Society of Nephrology : CJASN 9, 1974–1986, doi:10.2215/CJN.04480413 (2014).

22 Olinger, E., Schwaller, B., Loffing, J., Gailly, P. & Devuyst, O. Parvalbumin: calcium and magnesium buffering in the distal nephron. Nephrology, dialysis, transplantation : official publication of the European Dialysis and Transplant Association - European Renal Association 27, 3988–3994, doi:10.1093/ndt/gfs457 (2012).

23 Zaika, O., Tomilin, V., Mamenko, M., Bhalla, V. & Pochynyuk, O. New perspective of ClCKb/2 Cl-channel physiology in the distal renal tubule. American journal of physiology. Renal physiology 310, F923–930, doi:10.1152/ajprenal.00577.2015 (2016).

24 Roy, A., Al-bataineh, M. M. & Pastor-Soler, N. M. Collecting duct intercalated cell function and regulation. Clinical journal of the American Society of Nephrology : CJASN 10, 305–324, doi:10.2215/CJN.08880914 (2015).

25 Yang, Z. et al. Human Epididymis Protein 4: A Novel Biomarker for Lupus Nephritis and Chronic Kidney Disease in Systemic Lupus Erythematosus. Journal of clinical laboratory analysis 30, 897–904, doi:10.1002/jcla.21954 (2016).

26 Blomqvist, S. R. et al. Distal renal tubular acidosis in mice that lack the forkhead transcription factor Foxi1. The Journal of clinical investigation 113, 1560–1570, doi:10.1172/JCI20665 (2004).

27 Petrovic, S. et al. SLC26A7: a basolateral Cl-/HCO3-exchanger specific to intercalated cells of the outer medullary collecting duct. American journal of physiology. Renal physiology 286, F161–169, doi:10.1152/ajprenal.00219.2003 (2004).

28 Ohshiro, K. et al. Expression and immunolocalization of AQP6 in intercalated cells of the rat kidney collecting duct. Archives of histology and cytology 64, 329–338 (2001).

29 Chen, L. et al. Transcriptomes of major renal collecting duct cell types in mouse identified by single-cell RNA-seq. Proceedings of the National Academy of Sciences of the United States of America 114, E9989–E9998, doi:10.1073/pnas.1710964114 (2017).

30 Topper, J. N. et al. Human prostaglandin transporter gene (hPGT) is regulated by fluid mechanical stimuli in cultured endothelial cells and expressed in vascular endothelium in vivo. Circulation 98, 2396–2403 (1998).

31 Delprat, B. et al. FXYD6 is a novel regulator of Na,K-ATPase expressed in the inner ear. The Journal of biological chemistry 282, 7450–7456, doi:10.1074/jbc.M609872200 (2007).

32 Morita, K., Sasaki, H., Furuse, M. & Tsukita, S. Endothelial claudin: claudin-5/TMVCF constitutes tight junction strands in endothelial cells. The Journal of cell biology 147, 185–194 (1999).

33 Koda, R. et al. Novel expression of claudin-5 in glomerular podocytes. Cell and tissue research 343, 637–648, doi:10.1007/s00441-010-1117-y (2011).

34 Bondjers, C. et al. Transcription profiling of platelet-derived growth factor-B-deficient mouse embryos identifies RGS5 as a novel marker for pericytes and vascular smooth muscle cells. The American journal of pathology 162, 721–729, doi:10.1016/S0002-9440(10)63868-0(2003).

35 Stockand, J. D. & Sansom, S. C. Glomerular mesangial cells: electrophysiology and regulation of contraction. Physiological reviews 78, 723–744 (1998).

36 Smith, J. et al. Genes expressed by both mesangial cells and bone marrow-derived cells underlie genetic susceptibility to crescentic glomerulonephritis in the rat. Journal of the American Society of Nephrology : JASN 18, 1816–1823, doi:10.1681/ASN.2006070733(2007).

37 Segerer, S. et al. Compartment specific expression of dendritic cell markers in human glomerulonephritis. Kidney international 74, 37–46, doi:10.1038/ki.2008.99 (2008).

38 Stamatiades, E. G. et al. Immune Monitoring of Trans-endothelial Transport by Kidney-Resident Macrophages. Cell 166, 991–1003, doi:10.1016/j.cell.2016.06.058 (2016).

39 Villani, A. C. et al. Single-cell RNA-seq reveals new types of human blood dendritic cells, monocytes, and progenitors. Science 356, doi:10.1126/science.aah4573 (2017).

40 Watanabe, R. et al. Human skin is protected by four functionally and phenotypically discrete populations of resident and recirculating memory T cells. Science translational medicine 7,279ra239, doi:10.1126/scitranslmed.3010302 (2015).

41 Buggins, A. G. et al. Peripheral blood but not tissue dendritic cells express CD52 and are depleted by treatment with alemtuzumab. Blood 100, 1715–1720 (2002).

42 Lavin, Y. et al. Tissue-resident macrophage enhancer landscapes are shaped by the local microenvironment. Cell 159, 1312–1326, doi:10.1016/j.cell.2014.11.018 (2014).

43 Armulik, A., Genove, G. & Betsholtz, C. Pericytes: developmental, physiological, and pathological perspectives, problems, and promises. Developmental cell 21, 193–215, doi:10.1016/j.devcel.2011.07.001 (2011).

44 Esser, S. et al. Vascular endothelial growth factor induces endothelial fenestrations in vitro. The Journal of cell biology 140, 947–959 (1998).

45 Bekes, I. et al. Slit2/Robo4 Signaling: Potential Role of a VEGF-Antagonist Pathway to Regulate Luteal Permeability. Geburtshilfe und Frauenheilkunde 77, 73–80, doi:10.1055/s-0042-113461 (2017).

46 Yuen, D. A. & Robinson, L. A. Slit2-Robo signaling: a novel regulator of vascular injury. Current opinion in nephrology and hypertension 22, 445–451, doi:10.1097/MNH.0b013e32836235f4 (2013).

47 Chaturvedi, S. & Robinson, L. A. Slit2-Robo signaling in inflammation and kidney injury. Pediatric nephrology 30, 561–566, doi:10.1007/s00467-014-2825-4 (2015).

48 Liu, S. J. et al. Single-cell analysis of long non-coding RNAs in the developing human neocortex. Genome biology 17, 67, doi:10.1186/s13059-016-0932-1 (2016).

49 Hruz, T. et al. Genevestigator v3: a reference expression database for the meta-analysis of transcriptomes. Advances in bioinformatics 2008, 420747, doi:10.1155/2008/420747 (2008).

50 Stec, R., Grala, B., Maczewski, M., Bodnar, L. & Szczylik, C. Chromophobe renal cell cancer--review of the literature and potential methods of treating metastatic disease. Journal of experimental & clinical cancer research : CR 28, 134, doi:10.1186/1756-9966-28-134 (2009).

51 Kottgen, A. et al. Uromodulin levels associate with a common UMOD variant and risk for incident CKD. Journal of the American Society of Nephrology : JASN 21, 337–344, doi:10.1681/ASN.2009070725 (2010).

52 Coenen, M. J. et al. Phospholipase A2 receptor (PLA2R1) sequence variants in idiopathic membranous nephropathy. Journal of the American Society of Nephrology : JASN 24, 677–683, doi:10.1681/ASN.2012070730 (2013).

53 Liu, Q. Q. et al. Decreased DACH1 expression in glomerulopathy is associated with disease progression and severity. Oncotarget 7, 86547–86560, doi:10.18632/oncotarget.13470 (2016).

54 Al-Mayouf, S. M. et al. Loss-of-function variant in DNASE1L3 causes a familial form of systemic lupus erythematosus. Nature genetics 43, 1186–1188, doi:10.1038/ng.975 (2011).

55 Lu, Y., Ye, Y., Yang, Q. & Shi, S. Single-cell RNA-sequence analysis of mouse glomerular mesangial cells uncovers mesangial cell essential genes. Kidney international 92, 504–513, doi:10.1016/j.kint.2017.01.016 (2017).

56 Lu, Y. et al. Genome-wide identification of genes essential for podocyte cytoskeletons based on single-cell RNA sequencing. Kidney international 92, 1119–1129, doi:10.1016/j.kint.2017.04.022 (2017).

57 Der, E. et al. Single cell RNA sequencing to dissect the molecular heterogeneity in lupus nephritis. JCI insight 2, doi:10.1172/jci.insight.93009 (2017).

58 Huang, L. Y., Stuart, C., Takeda, K., D'Agnillo, F. & Golding, B. Poly(I:C) Induces Human Lung Endothelial Barrier Dysfunction by Disrupting Tight Junction Expression of Claudin-5. PloS one 11, e0160875, doi:10.1371/journal.pone.0160875 (2016).

59 Watanabe, T. et al. Paracellular barrier and tight junction protein expression in the immortalized brain endothelial cell lines bEND.3, bEND.5 and mouse brain endothelial cell 4. Biological & pharmaceutical bulletin 36, 492–59 (2013).

60 Tian, R. et al. The effect of claudin-5 overexpression on the interactions of claudin-1 and -2 and barrier function in retinal cells. Current molecular medicine 14, 1226–1237 (2014).

61 Mehta, D. & Malik, A. B. Signaling mechanisms regulating endothelial permeability. Physiological reviews 86, 279–367, doi:10.1152/physrev.00012.2005 (2006).

62 Ronnstrand, L. Signal transduction via the stem cell factor receptor/c-Kit. Cellular and molecular life sciences : CMLS 61, 2535–2548, doi:10.1007/s00018-004-4189-6 (2004).

63 Kandasamy, K. et al. NetPath: a public resource of curated signal transduction pathways. Genome biology 11, R3, doi:10.1186/gb-2010-11-1-r3 (2010).

64 Frey, S. K. et al. Isoforms of retinol binding protein 4 (RBP4) are increased in chronic diseases of the kidney but not of the liver. Lipids in health and disease 7, 29, doi:10.1186/1476-511X-7-29 (2008).

65 Ozcakar, Z. B. et al. DNASE1L3 mutations in hypocomplementemic urticarial vasculitis syndrome. Arthritis and rheumatism 65, 2183–2189, doi:10.1002/art.38010 (2013).

66 Strober, W.,Kitani, A.,Fichtner-Feigl, S. & Fuss, I. J. The signaling function of the IL-13Ralpha2 receptor in the development of gastrointestinal fibrosis and cancer surveillance. Current molecular medicine 9, 740–750 (2009)

67 Chandriani, S. et al. Endogenously expressed IL-13Ralpha2 attenuates IL-13-mediated responses but does not activate signaling in human lung fibroblasts. Journal of immunology193, 111–119, doi:10.4049/jimmunol.1301761 (2014).

68 Neelisetty, S. et al. Renal fibrosis is not reduced by blocking transforming growth factor-beta signaling in matrix-producing interstitial cells. Kidney international 88, 503–514, doi:10.1038/ki.2015.51 (2015).

69 Graeber, T. G. & Eisenberg, D. Bioinformatic identification of potential autocrine signaling loops in cancers from gene expression profiles. Nature genetics 29, 295–300, doi:10.1038/ng755 (2001).

70 Chang, Q., Ornatsky, O. & Hedley, D. Staining of Frozen and Formalin-Fixed, Paraffin-Embedded Tissues with Metal-Labeled Antibodies for Imaging Mass Cytometry Analysis. Current protocols in cytometry 82, 12 47 11–12 47 18, doi:10.1002/cpcy.29 (2017).

